# Learning enhances representations of taste-guided decisions in the mouse gustatory insular cortex

**DOI:** 10.1101/2023.10.16.562605

**Authors:** Joshua F. Kogan, Alfredo Fontanini

## Abstract

Learning to discriminate overlapping gustatory stimuli that predict distinct outcomes – a feat known as discrimination learning – can mean the difference between ingesting a poison or a nutritive meal. Despite the obvious importance of this process, very little is known on the neural basis of taste discrimination learning. In other sensory modalities, this form of learning can be mediated by either sharpening of sensory representations, or enhanced ability of “decision-making” circuits to interpret sensory information. Given the dual role of the gustatory insular cortex (GC) in encoding both sensory and decision-related variables, this region represents an ideal site for investigating how neural activity changes as animals learn a novel taste discrimination.

Here we present results from experiments relying on two photon calcium imaging of GC neural activity in mice performing a taste-guided mixture discrimination task. The task allows for recording of neural activity before and after learning induced by training mice to discriminate increasingly similar pairs of taste mixtures. Single neuron and population analyses show a time-varying pattern of activity, with early sensory responses emerging after taste delivery and binary, choice encoding responses emerging later in the delay before a decision is made. Our results demonstrate that while both sensory and decision-related information is encoded by GC in the context of a taste mixture discrimination task, learning and improved performance are associated with a specific enhancement of decision-related responses.

## INTRODUCTION

Animals must be able to discriminate similar, but not identical, sensory stimuli that predict different outcomes to adapt and survive in natural environments. In taste, for instance, discriminating small differences between otherwise similar solutions can be a matter of life and death. The ability to discriminate partially overlapping sensory stimuli can be enhanced with practice, a phenomenon also known as discrimination learning. While discrimination learning has been extensively investigated in a variety of sensory systems, taste discrimination learning remains surprisingly understudied.

The neural mechanisms underlying discrimination learning have typically been investigated in the context of operant paradigms in which subjects are trained to distinguish similar stimuli associated with different rewarded actions^1-5^. Neural recordings from these paradigms suggest two mechanistic models for learning, a sharpening of sensory representations and an enhancement of decision-related activity reflecting an improved ability to interpret sensory evidence for guiding decisions. The first has been observed in primary sensory cortices^2,3,6,7^, while the second is prevalent in “higher order” cortical regions involved in decision-making^8-11^.

Inferring from the literature which of the two models described above may be at play in the case of taste discrimination learning is not straightforward. The well-established role of the gustatory insular cortex (GC) in processing chemosensory information^12-15^ may point at the sharpening of taste representations as a key sensory mechanism for taste learning. However, recent evidence showing behaviorally relevant, decision-related signals^11,15-17^ in GC leaves open the possibility that taste learning may enhance cognitive signals. Observing the latter would be particularly interesting because it would suggest that a sensory area can enhance its ability to interpret sensory information to guide choices.

To study GC plasticity in the context of discrimination learning we relied on a taste mixture two alternative choice task (2-AC)^17^. We chose to use taste mixtures for a variety of reasons. First, in natural settings gustatory stimuli manifest mostly as mixtures. Second, mixtures allow for a systematic control of the degree of overlap between stimuli, making it easier to vary task difficulty and compare discrimination performance before and after training animals on progressively more overlapping pairs of stimuli^3,18,19^. Third, while GC processing of taste mixtures has never been investigated in the context of a discrimination task, studies across various modalities have described two types of neural responses to mixtures: monotonic linear responses tracking the intensity of mixture components and binary, categorical responses which could correspond to either the predominant mixture component or a binary choice associated with a learned category boundary^8-11,20-23^. These patterns can be used to track possible changes in sensory representations or decision-related activity.

In this study we present results from two photon calcium imaging of GC in mice trained on a taste mixture 2-AC which utilized a delay period to separate sensation from choice. Mice were first trained to discriminate between pure sucrose and NaCl. They were then tested using sucrose/NaCl mixtures and asked to identify the predominant taste in the mixture. Discrimination performance dropped to near chance levels with highly overlapping taste mixtures. Analysis of neural activity revealed responses that significantly differed from those described in rats passively receiving taste mixtures^21^. Specifically, in the sampling period neural responses varied linearly with mixture concentration. In contrast, neural responses in the delay period were categorical and encoded binary responses to the upcoming choice. Mice were then trained to discriminate between progressively more similar mixtures, which led to improved behavioral performance. Comparison of the sensory and decision-related responses in the pre- and post-learning conditions revealed a selective enhancement of decision-related neural responses after learning.

We conclude that in a taste-guided mixture discrimination task, both sensory and decision-related information is encoded in GC neural activity. Further, upon developing a novel training paradigm for taste discrimination learning, we demonstrate that learning is associated with a specific enhancement of decision-related signals.

## RESULTS

### Taste Mixture Discrimination 2-AC Task

To investigate the involvement of GC in taste mixture discriminations, six head-fixed, water-restricted mice were trained to perform a two-alternative choice (2-AC) taste discrimination task. Mice learned to sample a taste stimulus from a central spout and respond by licking one of two lateral spouts. Correct choices were rewarded with a drop of water and errors punished with a timeout (**Fig 1a**). This task is designed to engage both sensory and cognitive processes, as mice need to first identify a gustatory stimulus (taste solution from the central spout), correctly associate that stimulus with an outcome (reward left vs right) and make a choice. A delay period (∼3.5 s) allows for temporal separation of the sensory and decision-related processes in the trial (**Fig 1b**).

**Figure 1.**
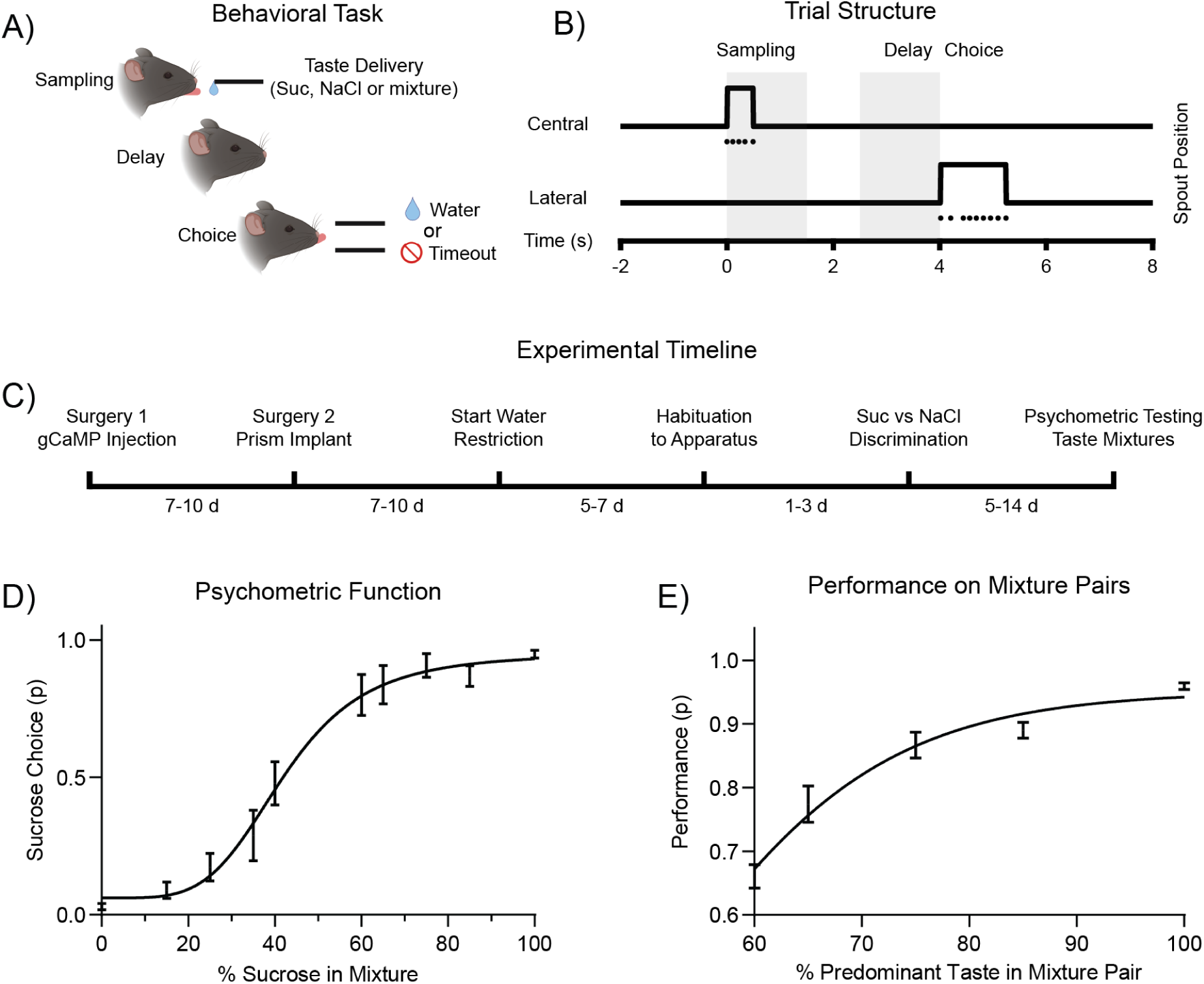
Taste Mixture Discrimination 2-AC Task. **A)** Schematic diagram of the 2-AC behavioral task. **B)** Schematic diagram showing the outline of the trial structure, with sampling, delay, and choice periods. Grey shading indicates 1.5 s long windows used for subsequent analysis of activity related to sampling and delay. Each dot represents one lick. **C)** Schematic diagram showing the experimental timeline for surgeries and behavioral training. **D)** Plot showing the mean behavioral performance for all the mixture stimuli (n = 6 mice). Error bars represent standard error of the mean (SEM). The black line represents sigmoid fit to the mean performance data (r^2^ = 0.99, IC50 = 43.77, slope = 0.038). **E)** Plot showing the mean behavioral performance for all mixture pairs (n = 6 mice). Error bars represent SEM. The black line represents the logistic fit to the mean performance data (r^2^ = 0.84, Y_max_ = 0.95, Y_0_ = 0, k = 0.10).

Mice were first trained to discriminate between pure sucrose (100 mM) and NaCl (100 mM). After reaching criterion performance on this part of the task (>85% correct trials for three consecutive days), mice were presented with taste mixtures across two testing sessions (%Sucrose/%NaCl in mixture, Day 1: 100/0, 85/15, 65/35, 35/65, 15/85, 0/100; Day 2: 100/0, 75/25, 60/40, 40/60, 25/75, 0/100) to obtain a psychometric function. Mice had only been trained on the pure sucrose vs NaCl discrimination and had not previously experienced any of the taste mixtures (**Fig 1c**). For these sessions, animals followed the rule of selecting the side associated with the predominant taste in the mixture to receive a water reward. We calculated average behavioral performance for each mixture and fit a sigmoidal curve to obtain a psychometric function (r^2^ = 0.99, IC50 = 43.77, slope = 0.038, **Fig 1d**). Calculating correct responses for mixture pairs showed that the performance deteriorates as the mixtures become closer to 50/50 (one-way ANOVA, F(4,25) = 38.22, p < 0.001, chance performance, 50%, **Fig 1e**). Analysis of licking for the ten different stimuli revealed no significant difference with varying mixture concentrations in sampling duration (one-way ANOVA, F(9,62) = 0.09, p = 0.99) or latency of lateral licking (one-way ANOVA, F(9,62) = 0.51, p = 0.86).

### Task Related Responses in GC

Two photon calcium imaging was used to record activity from the superficial layers of GC in mice engaged in the psychometric testing sessions (**Fig 2a-c**). Fluorescence signals were extracted from neurons expressing gCaMP7f, 2337 neurons were recorded from six mice (416.3 ± 19.4 neurons per mouse).

**Figure 2.**
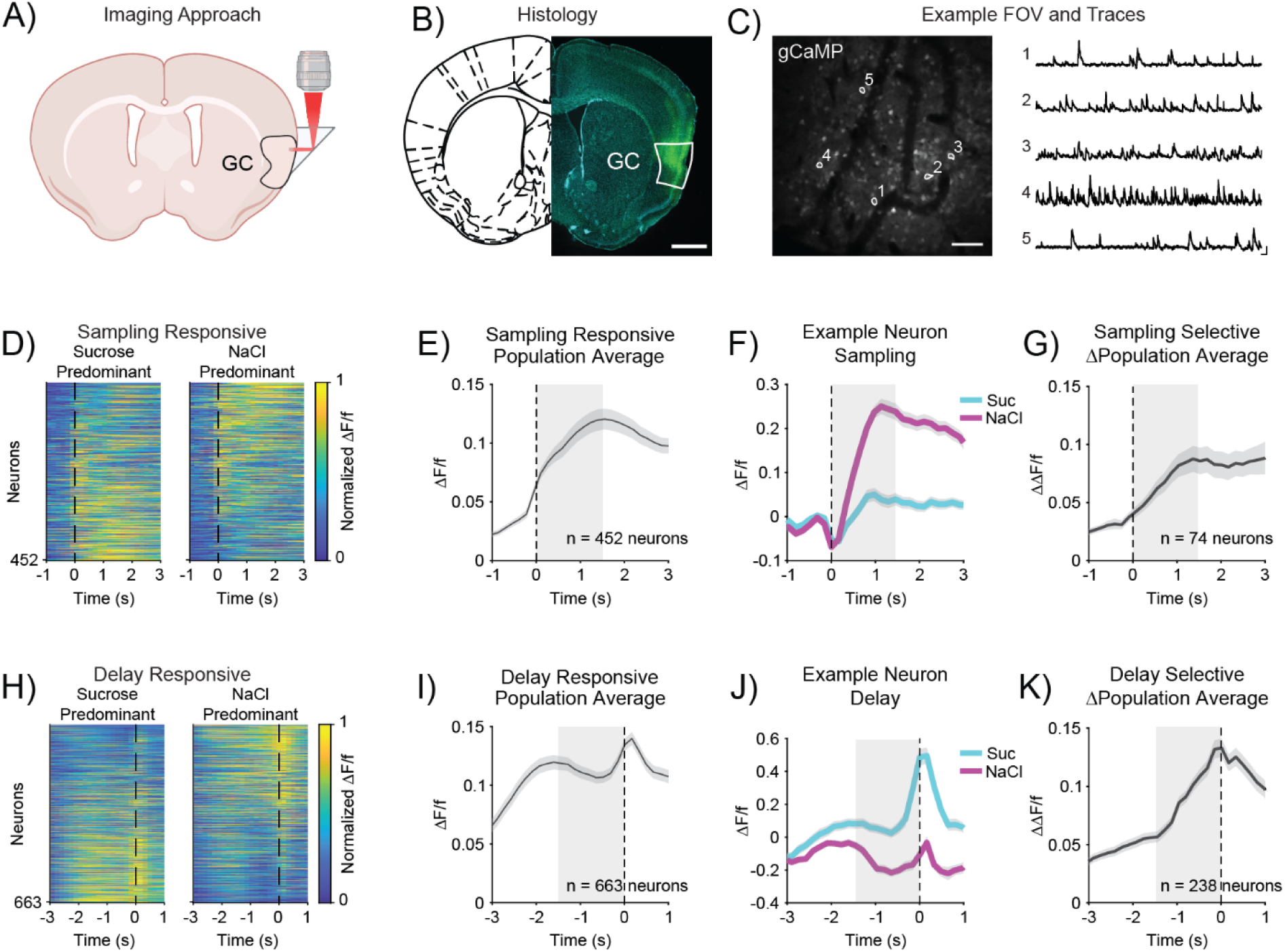
Task Related Responses in GC. **A)** Schematic diagram showing the microprism-based approach for two photon calcium imaging in GC. **B)** Image of a representative coronal section showing gCaMP expression (green) in GC and Hoechst counterstain (cyan). The scale bar represents 1 mm. **C)** Left. Image of an example field of view (FOV) with selected regions of interest (ROIs), scale bar represents 100 µm. Right. Representative ΔF/f traces corresponding to ROIs shown on left. The horizontal scale bar represents 13 s and the vertical scale bar represents a ΔF/f value of 0.5. **D)** Heatmaps showing sampling responsive neurons, aligned to onset of taste sampling (t = 0 s). Activity is normalized across both trial types and sorted by difference in activity between trial types in the sampling period (1.5 s after sampling onset). **E)** Plot showing population average response of sampling responsive neurons across all trials (same as neurons in D, n = 452 neurons). **F)** Plot showing example sampling responsive neuron. Sampling onset occurs at t = 0 s. **G)** Plot of delta population average showing the mean absolute difference in activity between sucrose and NaCl trials for responsive neurons that are trial type selective in the sampling period (74/452 neurons). **H)** Heatmaps showing delay responsive neurons, aligned to onset of choice (t = 0 s). Activity is normalized across both trial types and sorted by difference in activity between trial types in the delay period (1.5 s before choice). **I)** Plot showing population average response of delay responsive neurons across all trials (same as neurons in H, n = 663 neurons). **J)** Plot showing example delay responsive neuron. Choice occurs at t = 0 s. **K)** Plot of delta population average showing the mean absolute difference in activity between sucrose and NaCl trials for delay responsive neurons that are trial type selective in the delay period (238/663 neurons). For E-G and I-K, shading indicates SEM and grey box indicates analysis window.

To begin assessing task responsiveness, single neuron activity during the sampling and delay periods was compared to baseline. Consistent with previous studies in GC employing a 2-AC^17^, we observed significant modulations of activity during the task epochs. Both sucrose- and NaCl-predominant trials showed modulations compared to baseline (**Fig 2d-e, h-i**), with 19.3% [452/2337] of neurons responding within 1.5 s from taste delivery (“sampling responsive”) and 28.4% [663/2337] responding in the delay period, during a 1.5 s window just before choice (“delay responsive”). A subset of task-responsive neurons showed trial type selectivity with significantly different responses to sucrose- vs NaCl-predominant trials. Specifically, 16.4% [74/452] of neurons were selective during the sampling period and 35.4% [238/663] during the delay. **Figure 2f,j** show representative examples of neurons selective to trial type (i.e., sucrose- vs NaCl-predominant).

To assess the time course of selectivity, the absolute difference between trial types was computed for selective neurons and averaged across neurons for the sampling and delay periods (Δ population average, **Fig 2g,k**). **Figure 2g,k** shows that the difference in activity for neurons selective during the sampling period peaks 1.4 s after taste delivery (maximum ΔΔF/f = 0.086), while the difference for neurons selective during the delay period peaks at 0.15 s before the lateral lick (maximum ΔΔF/f = 0.132).

In summary, calcium imaging experiments show that a significant portion of GC neurons are task-responsive and differentiate trial types during the sampling and delay periods.

### Linear and Binary Response Patterns

Trial type selectivity could result from the encoding of either the intensity of one or both components of the taste mixtures, the predominant taste quality (sucrose vs NaCl) or upcoming choice (left vs right). Encoding of component intensity of mixtures is typically linear^8,11,21,24^, while encoding of categorical taste quality or upcoming choice is binary^8-11^. To assess whether coding of the various stimuli was linear or binary, we fit single neurons responses to the individual taste mixtures with both linear and sigmoid curves (example responses and fits, **Fig 3a-b**). Neural responses were binned into 0.5 s bins and analyzed in both the sampling and the delay periods. F values were compared to assess goodness of fit and a best fit type was calculated for each neuron in each bin. This approach accounts for the higher number of parameters required to fit the sigmoid compared to the linear function^21^ (see methods for details). We observed neurons whose activity could be fit with either linear or sigmoid functions. Neurons with a best fit to the linear pattern could be observed in both the sampling and the delay periods, consistent with previously observed taste mixture coding (sampling period linear fits: 9% [41/452], maximum fits 1 s after sampling onset; delay period linear fits: 18% [122/663], maximum fits 0.5 s before choice, chance level for fits = 5%, **Fig 3c-d**). In contrast, neurons following the sigmoid fit pattern only emerged in the delay (sampling period sigmoid fits: 2% [9/452]; delay period sigmoid fits: 11% [74/663], maximum fits 0.5 s before choice, chance level for fits = 5%). There was also an overall increase in both types of fits when comparing the sampling with the delay periods (linear fits, χ^2^ test, p < 0.001; sigmoid fits, χ^2^ test, p < 0.001, **Fig 3d**). These results suggest neural activity encoding mixture identity exists throughout the sampling and delay periods, while binary responses to the upcoming choice emerge only in the delay.

**Figure 3.**
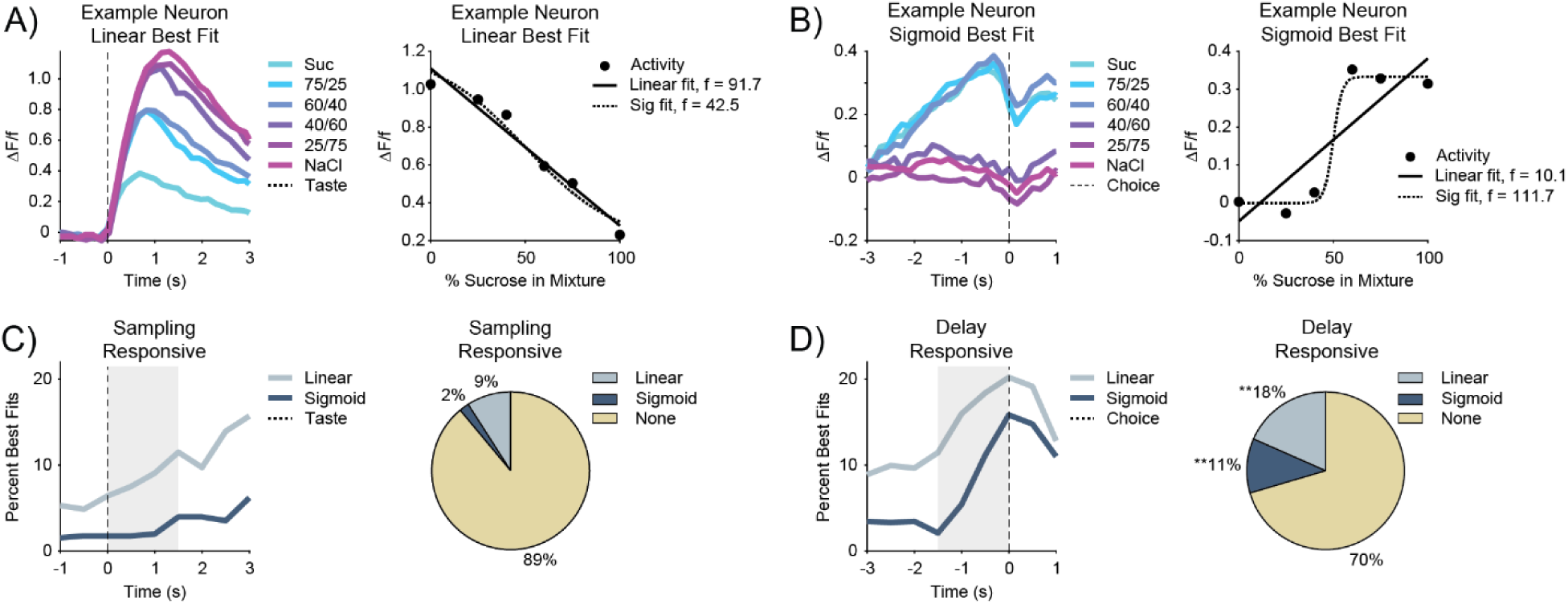
Linear and Binary Response Patterns. **A)** Left. Plot showing a representative neural response to different sucrose and NaCl mixtures (Suc/NaCl expressed in %) with a linear best fit, taste delivery occurs at t = 0 s. Right. Plot showing example linear (solid line, f = 91.7) and sigmoid (dashed line, f = 42.5) fits to the response on left. **B)** Left. Plot showing a representative neural response to different sucrose and NaCl mixtures (Suc/NaCl expressed in %) with a sigmoid best fit, choice occurs at t = 0 s. Right. Plot showing example linear (solid line, f = 10.1) and sigmoid (dashed line, f = 111.7) fits to the response on left. **C)** Left. Plot showing the proportion of sampling responsive neurons with best fit to linear or sigmoid patterns, taste delivery occurs at t = 0 s. Right. Pie chart showing the maximum proportion of fit types in the sampling period (1.5 s after taste delivery). **D)** Left. Plot showing the proportion of delay responsive neurons with best fit to linear or sigmoid patterns, choice occurs at t = 0 s. Right. Pie chart showing the maximum proportion of fit types in the delay epoch (1.5 s before choice). Asterisk indicates higher proportion of fits in delay versus sampling period, Χ² test, **p<0.001.

Next, mixture responses were analyzed at the population level using a maximum correlation decoder^25^ (see methods for details). Neurons from all animals (n = 6, 2337 neurons) were combined to generate a pseudo-population, which was used for all subsequent analyses. Individual decoders were trained using correct trials from each mixture pair. When trained on activity from the sampling period, decoding performance for the pure tastes was close to 100%, dropping to near chance as the stimuli become more similar (100/0 vs 0/100: 93.9%, 75/25 vs 25/75: 78.6%, 60/40 vs 40/60: 53.0%: chance 50%, **Fig 4a, circles**). In contrast, activity from the delay period was able to decode all mixture pairs close to 100% (100/0 vs 0/100: 99.8%, 75/25 vs 25/75: 99.6%, 60/40 vs 40/60: 88.1%: chance 50%, **Fig 4a**,

**Figure 4.**
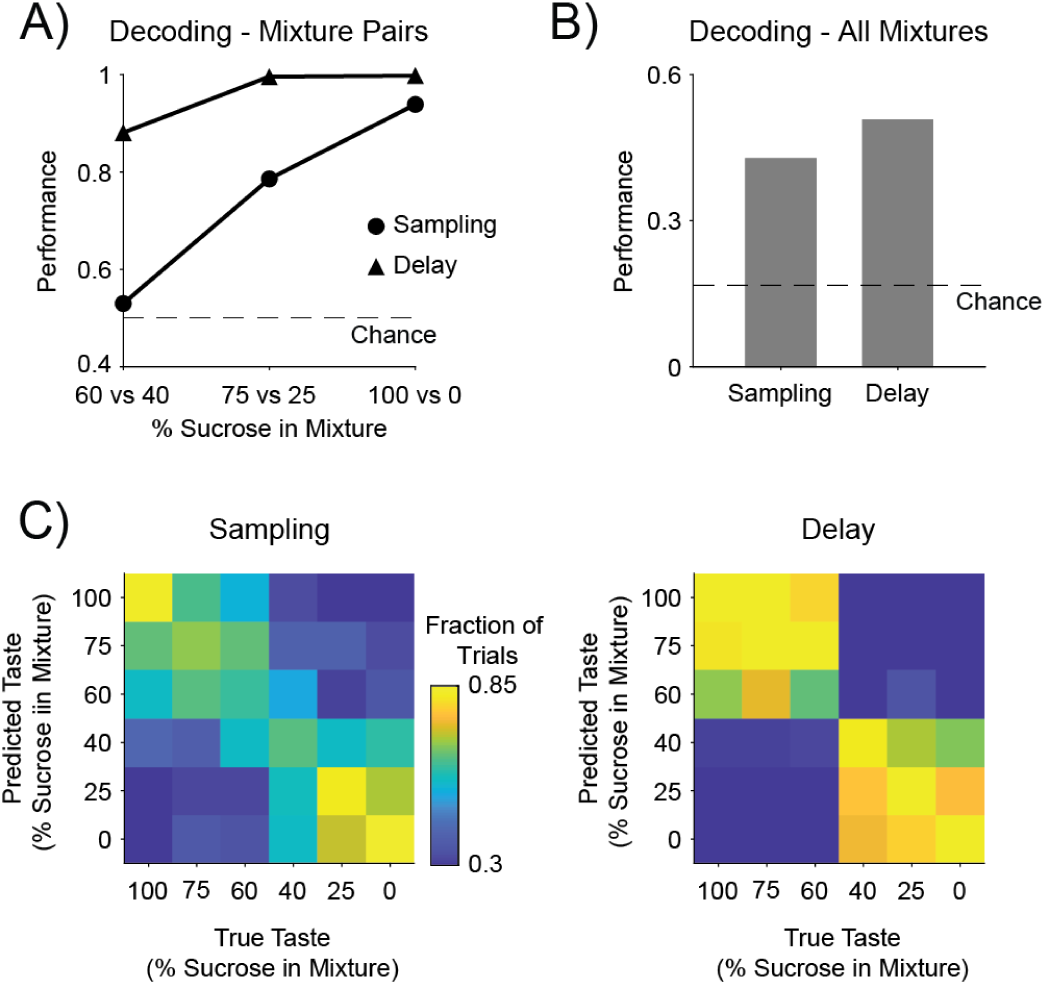
Population Decoding of Mixture Responses. **A)** Plot showing the decoding performance for each mixture pair in the sampling and delay periods, value on X axis indicates the percentage of sucrose in the mixture. Dashed line indicates chance level performance. **B)** Plot showing decoding performance for all mixtures (n = 6) in the sampling and delay periods. Dashed line indicates chance level performance. **C)** Heatmap plots showing the confusion matrices for decoding all mixtures in the sampling and delay periods (same decoders as B). Labels indicate the percentage of sucrose in the mixture.

**triangles**). To assess overall decoding of mixture identity, decoders were trained on all six trial types. There was similar overall decoding accuracy in both task epochs, with a slight enhancement in the delay (sampling: 42.8%, delay: 50.7%, chance: 16.6%, **Fig 4b**). To better evaluate decoding performance, confusion matrices were generated. Consistent with the decoders trained on individual mixture pairs, activity in the sampling period showed high performance for the pure tastes with a graded decrease for increasingly similar pairs (**Fig 4c, left**), while delay period activity showed high confusion for all mixture pairs that indicated the same direction, and low confusion for pairs that indicate the opposite direction (**Fig 4c, right**).

Altogether, these results demonstrate that GC activity can represent mixtures linearly as well as binarily, and that binary representations (and decoding) are more pronounced in the delay period preceding lateral licking.

### Delay Period Activity and Coding of Upcoming Choice

To disambiguate whether the observed binary delay activity encoded the prevailing taste category (sucrose- vs NaCl-predominant) or the upcoming choice (left vs right), error trials were analyzed. Stimulus and choice selectivity indexes were calculated for each of the sampling and delay responsive neurons. The stimulus selectivity index represents the difference between sucrose and NaCl responses on correct trials, while choice selectivity calculates the mean difference between responses in correct and error trials where the stimulus was the same, but the choice was different. Two example neurons demonstrate the difference between a stimulus selective neuron that is not choice selective (**Fig 5a**) and one that is choice selective (**Fig 5c**). Briefly, the neuron that is exclusively stimulus selective (and has no choice selectivity) displays the same pattern of taste responsiveness in correct and error trials (sucrose > NaCl). On the contrary, the stimulus selective neuron that is also choice selective shows opposite taste responses in correct and error trials (correct: NaCl > sucrose; error: sucrose > NaCl); what remains constant across correct and error trials is the choice selectivity (right > left). This neuron’s activity represents planned licking direction. Plotting stimulus vs choice selectivity for all responsive neurons showed a stronger correlation in the delay vs the sampling periods (sampling: r^2^ = 0.099, slope = 0.428, p < 0.001; delay: r^2^ = 0.397, slope = 0.666, p < 0.001, **Fig 5b,d**), indicating that the delay period contains information related to the upcoming choice.

**Figure 5.**
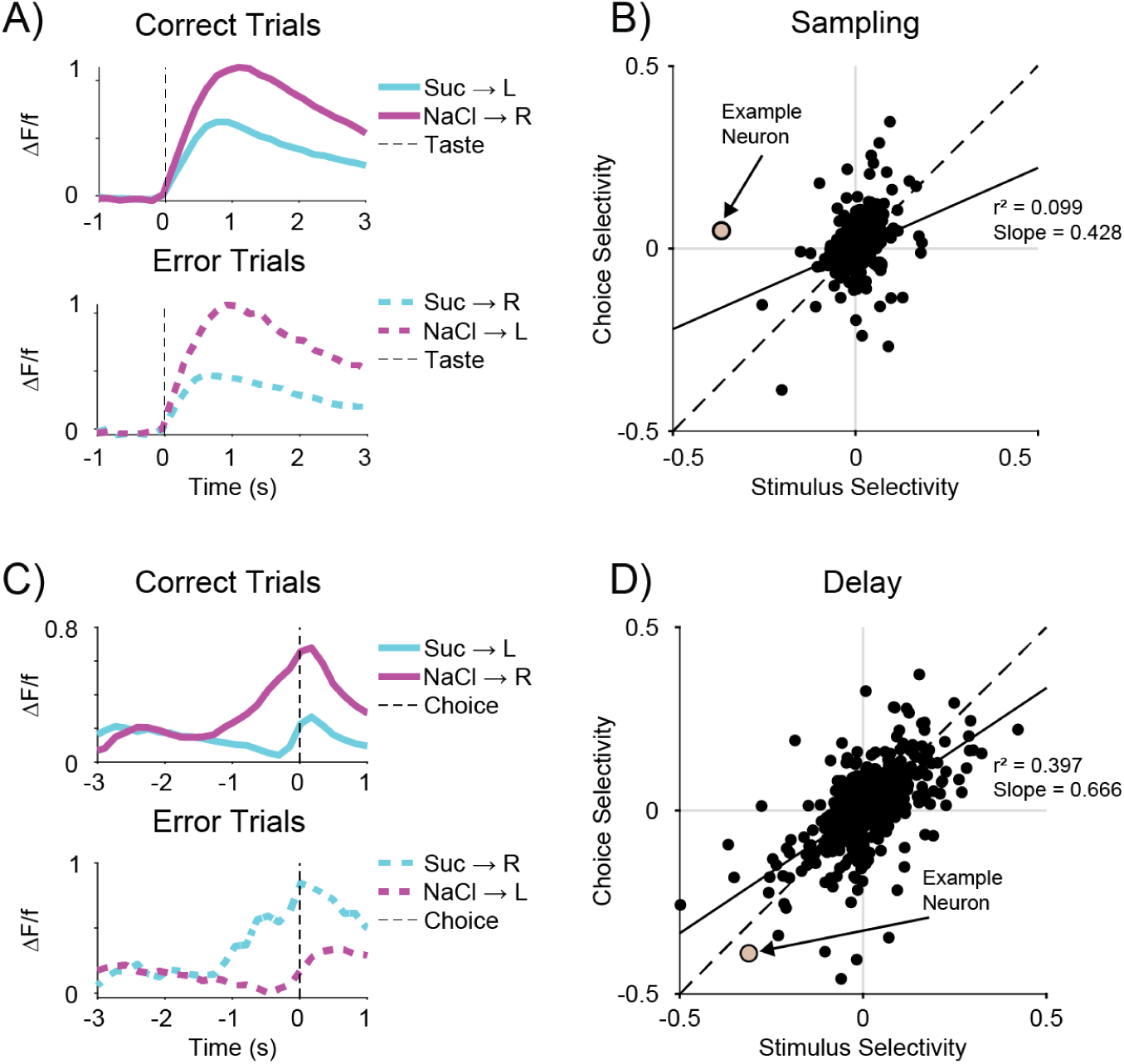
Delay Period Activity and Coding of Upcoming Choice. **A)** Plot for one example neuron showing the average responses to sucrose and NaCl for correct (top) and error (bottom) trials. This neuron is selective to the stimulus regardless of the choice made by the animal. Taste delivery occurs at t = 0 s. **B)** Scatter plot showing correlation between stimulus and choice selectivity for each responsive neuron in the sampling window (1.5 s after taste delivery). Example neuron from A is labeled on the plot. The solid line represents a linear fit to the data (r^2^ = 0.099, slope = 0.428, p < 0.001) and the dashed line represents the unity line. **C)** Plot for one example neuron showing the average responses to sucrose and NaCl for correct (top) and error (bottom) trials. This neuron is selective to the choice rather than the stimulus identity. Choice occurs at t = 0 s. **D)** Scatter plot showing correlation between stimulus and choice selectivity for each responsive neuron in the delay window (1.5 s before choice). Example neuron from C is labeled on the plot. The solid line represents a linear fit to the data (r^2^ = 0.397, slope = 0.666, p < 0.001) and the dashed line represents the unity line.

### Improvement of Behavioral Performance After Discrimination Learning

Upon establishing a mixture discrimination paradigm and identifying the associated patterns of activity, we elected to investigate whether behavioral performance and GC neural activity could be modified by training mice in a novel, taste mixture discrimination learning paradigm. Mice trained on the initial sucrose (100 mM) vs NaCl (100 mM) paradigm and tested with the battery of mixtures were further trained to discriminate pairs of progressively more similar sucrose/NaCl mixtures (75/25 vs 25/75; 65/35 vs 35/65; 60/40 vs 40/60; **Fig 6a**). After reaching criteria performance (>80% correct choices on the 60/40 vs 40/60 pair), they were tested on post-learning psychometric sessions, with the same panel of mixtures as the pre-learning sessions. A sigmoid curve was fit to the post-learning data to obtain a psychometric function (r^2^ = 0.99, IC50 = 52.14, slope = 0.055, **Fig 6b**, green). The pre- and post-learning psychometric curves were better fit by two curves than one (n = 6 animals, extra-sum-of-squares F test, F(4,112) = 9.9626, p < 0.001, **Fig 6b**) and performance was higher for all mixture pairs after learning such that the data were better fit by two logistic curves than one (pre: r^2^ = 0.84, Y_max_ = 0.95, Y_0_ = 0, k = 0.10, post: r^2^ = 0.89, Y_max_ = 0.99, Y_0_ = 0, k = 0.15, extra-sum-of-squares F test, F(4,54) = 24.57, p < 0.001, **Fig 6c**). These results demonstrate that specific pair training enhances mice’s ability to discriminate and categorize a panel of mixtures.

**Figure 6.**
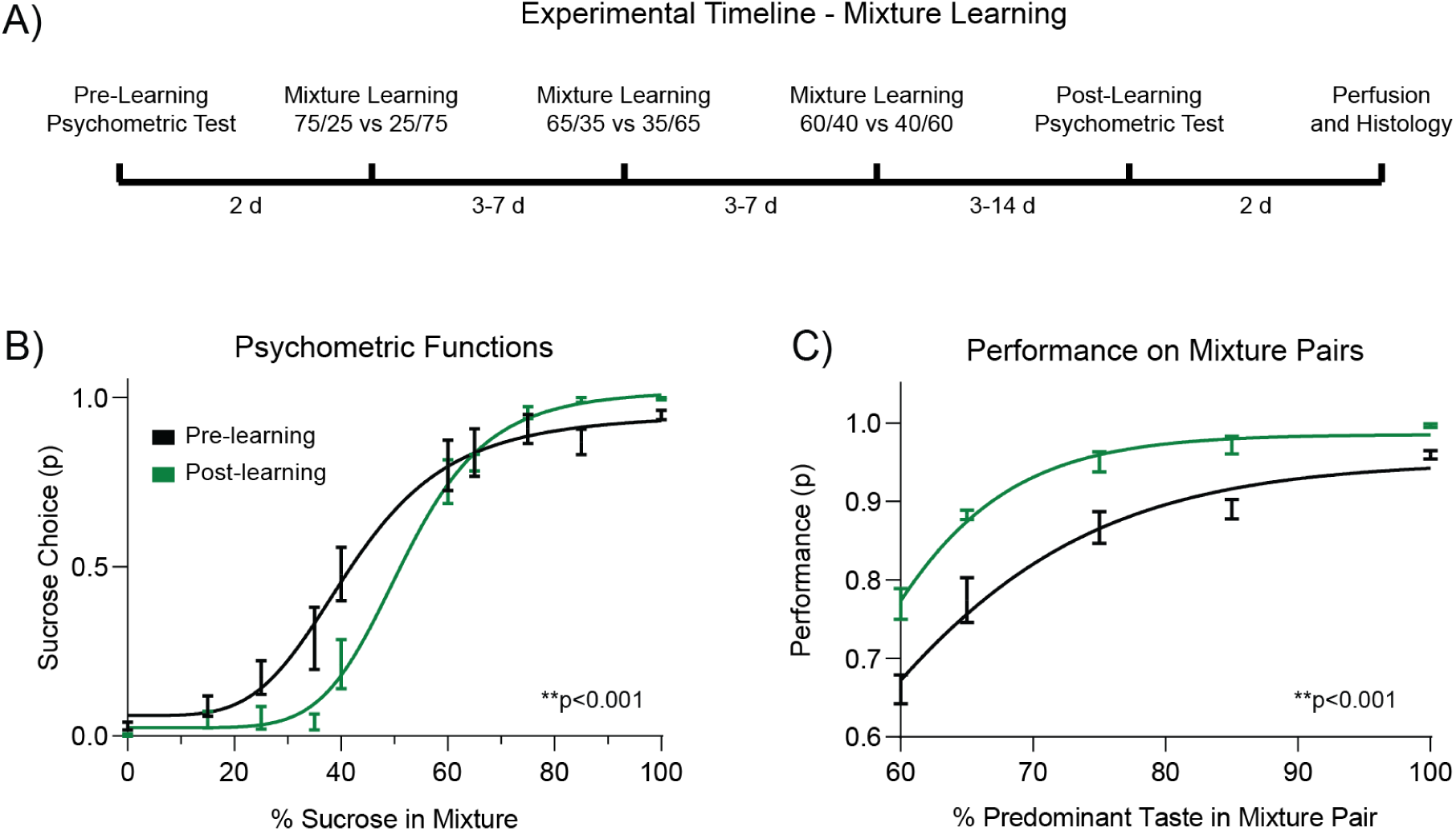
Improvement of Behavioral Performance After Discrimination Learning. **A)** Schematic diagram showing the experimental timeline for the mixture discrimination training. **B)** Plot showing the mean behavioral performance for all the mixture stimuli in the pre- and post-learning conditions (n = 6 mice). Error bars represent SEM. Black line represents sigmoid fit to the mean pre-learning performance data (same as Fig 1b, r^2^ = 0.99, IC50 = 43.77, slope = 0.038). Green line represents sigmoid fit to the mean post-learning performance data (r^2^ = 0.99, IC50 = 52.14, slope = 0.055). The data is better fit by two curves than one, extra-sum-of-squares F test, **p < 0.001. **C)** Plot showing the mean behavioral performance for all the mixture pairs in the pre- and post-learning conditions (n = 6 mice). Error bars represent SEM. Black line represents logistic fit to the mean pre-learning performance data (same as Fig 1c, r^2^ = 0.84, Y_max_ = 0.95, Y_0_ = 0, k = 0.10). Green line represents fit to the mean post-learning performance data (r^2^ = 0.89, Y_max_ = 0.99, Y_0_ = 0, k = 0.15). The data is better fit by two curves than one, extra-sum-of-squares F test, **p < 0.001.

### Mixture Discrimination Learning and Enhanced Decision-Related Activity

Neural responses were compared between pre- and post-training sessions to identify changes that occurred across discrimination learning. There was no significant difference in the proportion of responsive neurons in either the sampling or delay period (sampling, pre: 19.3% [452/2337], post: 18.6% [493/2658], pre vs post, χ^2^ test, p > 0.05; delay, pre: 28.4% [663/2337], post: 28.5% [758/2658], χ^2^ test, p > 0.05, **Fig 7a**). However, there was a significant difference in the proportion of neurons showing trial type selectivity after learning, with a pronounced increase in the delay period (sampling, pre: 16.4% [74/452], post: 11.4% [56/493], pre vs post, χ^2^ test, p < 0.05; delay, pre: 35.4% [235/663], post: 54.1% [410/758], χ^2^ test, p < 0.0001, **Fig 7b**). There was also an overall increase in the selectivity of the delay responsive neurons (sampling selectivity pre: 0.028, post: 0.025, unpaired t test, p = 0.23; delay selectivity pre: 0.059, post: 0.077, unpaired t test, p < 0.001, **Fig 7c**) and an earlier onset of delay selectivity in the pre- vs post-learning conditions (half maximum selectivity; pre: 0.47 s before choice, post: 0.79 s before choice, **Fig 7d**).

**Figure 7.**
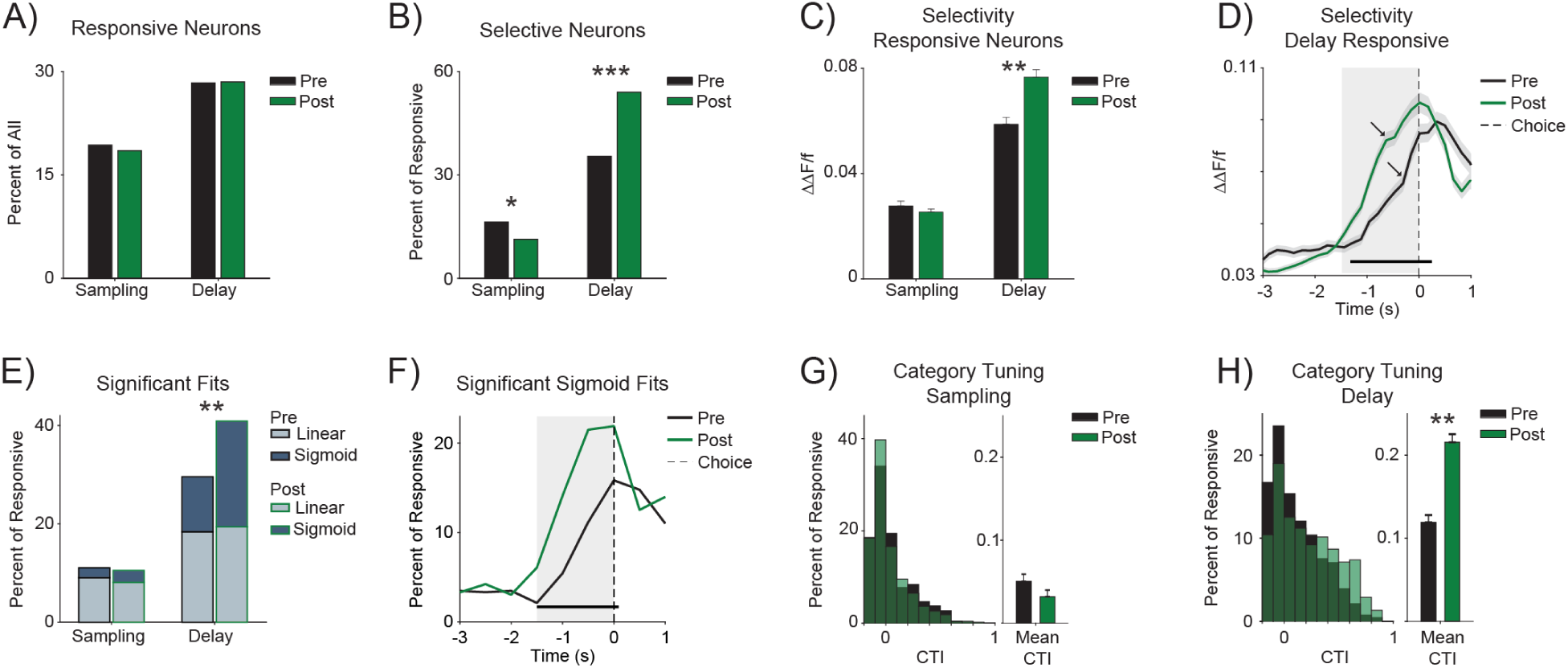
Mixture Discrimination Learning and Enhanced Decision-Related Activity. **A)** Plot showing the proportions of sampling and delay responsive neurons in pre- and post-learning conditions. **B)** Plot showing the proportions of responsive neurons with selectivity to taste mixture categories. Asterisk indicates significant difference between pre- and post-learning conditions, χ² test, *p < 0.05, ***p<0.0001. **C)** Plot showing mean selectivity of sampling and delay responsive neurons (correct trials only). Asterisk indicates significant difference between pre- and post-learning conditions, unpaired t test, **p<0.001. **D)** Plot showing the time course of selectivity for delay responsive neurons pre- and post-learning. Choice occurs at t = 0 s and grey box indicates delay window (1.5 s before choice). Shading indicates SEM. Horizontal black line indicates a significant difference between the two conditions, unpaired t test, p < 0.05, Bonferroni corrected. Time of half max selectivity in the delay window (arrows), pre = 0.47 s before choice, post = 0.79 s before choice. **E)** Plot showing the percent of responsive neurons with a best linear or sigmoid fit to trial averaged mixture responses in the sampling and delay windows. Asterisk indicates significant difference between pre- and post-learning conditions, χ² test, **p < 0.001. **F)** Plot showing the time course of the proportion of responsive neurons with best sigmoid fits in the pre- and post-learning conditions. Choice occurs at t = 0 s and grey box indicates delay window (1.5 s before choice). Horizontal black line indicates a significant difference between the two conditions, χ² test, p < 0.05, Bonferroni corrected. **G)** Plot showing the category tuning index (CTI) distributions (left) and mean (right) for pre- and post-learning sampling responsive neurons in the sampling window. **H)** Plot showing the category tuning index (CTI) distributions (left) and mean (right) for pre- and post-learning delay responsive neurons in the delay window. Asterisk indicates significant difference between pre- and post-learning conditions, unpaired t test, **p<0.001. For all plots, pre-learning data is shown in black and post-learning data is shown in green.

Curve fitting analysis comparing pre- vs post-learning mixture responses in the delay period showed no change in the proportion of linear fits, but a significant increase in neurons with a best fit for the sigmoid pattern (linear pattern, pre: 18.4% [122/663], post: 19.4% [147/758], χ^2^ test, p = 0.184; sigmoid pattern, pre: 11.6% [74/663], post: 21.5% [163/758], χ^2^ test, p < 0.001, **Fig 7e**). After learning, examination of the time course of sigmoid fits in the delay revealed a significantly higher proportion of neurons displaying this pattern in all time bins (1.5 s to 1 s before choice, pre: 2.2% [14/663], post: 6.1% [46/758]; 1 s to 0.5 s before choice, pre: 5.4% [36/663], post: 14.1%, [107/758]; 0.5 s to 0 s before choice, pre 11.2% [74/663], post: 21.5% [163/758]; χ^2^ test for all bins, p < 0.0001, **Fig 7f**). These data suggest that discrimination learning is associated with enhanced categorical responses in the delay period that are selective to the upcoming decision. Like for the pre-learning condition, analysis of error trials in the post-learning condition showed a stronger correlation between stimulus and choice selectivity in the delay vs sampling post-learning (sampling: r^2^ = 0.073, slope = 0.548, p < 0.001; delay: r^2^ = 0.426, slope = 0.596, p < 0.001, **Supplemental Fig 1**), suggesting this selectivity is related to the upcoming choice, rather than a categorical perceptual representation of the mixture stimuli.

Lastly, a category tuning index (CTI) was calculated for all sampling and delay responsive neurons^9,10^. CTI measures the normalized difference between taste stimuli that indicate the same choice and those that indicate opposite choices. This value will be larger if responses are similar across all stimuli that indicate the same choice, and different from those that indicate the opposite choice. There was no difference in the sampling CTI, but a significant increase in the delay CTI after learning (sampling: mean CTI pre: 0.05, post: 0.03, unpaired t test, p = 0.12; delay: mean CTI pre: 0.12, post: 0.22, unpaired t test, p < 0.001 **Fig 7g-h**). This finding is also consistent with the enhancement of categorical decision-related representations in the delay period after learning.

Overall, these results show that taste discrimination training leading to improved behavioral performance is associated with enhanced decision-related responses rather than changes in the stimulus representation.

## DISCUSSION

The experiments presented in this manuscript rely on two photon calcium imaging to: i) unveil patterns of neural activity representing taste mixtures and decision-related information in mice engaged in a taste discrimination task; and ii) demonstrate that decision-related signals can be enhanced by training mice in a discrimination learning paradigm.

To study taste mixture discrimination we relied on a 2-AC task^17,19,26,27^. Mice were first trained to associate sucrose and NaCl delivered at a central spout with a water reward delivered at either a left or right lateral spout. Upon learning the basic sucrose vs NaCl discrimination, mice were tested on a mixture discrimination, where they were presented with varying mixtures of sucrose and NaCl, and were rewarded for selecting the direction associated with the predominant taste in the mixture. The task included a delay period separating the sampling and choice, which allowed for the examination of the temporal evolution of neural activity as animals engage with each trial of the task. Two photon calcium imaging of GC’s superficial layers showed modulations of neural activity during both the sampling and delay periods. To further elucidate the nature of these responses, we analyzed single neuron responses by fitting tuning curves. Consistent with the existing literature on mixture coding^21^, linear fitting was used to assess the linear encoding of mixture element concentrations. Sigmoid fitting was used to extract responses that encoded for binary categories. We show that neurons display linear responses to the mixtures as early as the sampling period. On the contrary, categorical responses emerge in the delay period, where neurons display binary responses to the upcoming choice, regardless of which mixture was presented on that trial. Analysis of error trials confirmed that categorical responses were related to the upcoming choice (left vs right) and not simply to the general taste quality (sweet vs salty).

To investigate the plasticity of sensory and decision-related signals we trained mice in a multi-day discrimination learning paradigm using progressively more overlapping pairs of mixtures. Mixture discrimination testing, using the same 2-AC paradigm adopted before training, was performed after learning to demonstrate that mice can learn and improve their taste discrimination performance. This behavioral paradigm was inspired by protocols adopted for perceptual learning in other sensory systems ^2,5,8,18,19,28^. In addition to being the first to demonstrate taste discrimination learning in the context of a reward-directed operant task, this paradigm is designed to monitor neural activity before and after learning. Imaging during the testing sessions revealed a specific increase in categorical responses in the delay period, consistent with an improvement in the ability to interpret sensory signals to enhance discrimination performance.

Altogether the results presented here demonstrate that GC multiplexes processing of sensory and decision-related signals and that the latter can be enhanced by learning.

### Taste Mixture Coding

In natural settings, animals seldom consume pure tastants, in fact they typically experience mixtures of stimuli^20^. Only a handful of studies have investigated the coding of mixtures in the gustatory system of alert animals^21,29,30^. Electrophysiological experiments on the GC of rats passively receiving taste mixtures via an intraoral cannula explored three patterns of possible responses: i) monotonic responses largely dependent on the concentration of an individual component; ii) categorical responses tracking the quality of the prevailing element; and iii) non-linear responses reflecting mixture enhancement or suppression^21^. Analysis of the time course of firing responses provided evidence for early and brief mixture suppression followed by a prevalence of linear monotonic responses. Interestingly, no evidence was found for categorical coding of mixture stimuli, which has been observed in other sensory systems^22^.

In the present study, we used mixtures of 100 mM sucrose and 100 mM NaCl, linearly varying the concentrations of each of the components. In agreement with previous studies, we found that representations of mixtures varied during the time course of a single trial, and that the majority of single neuron responses linearly represented mixtures. Other patterns of responsiveness, however, differed from those reported in passively tasting rats. While we did observe isolated responses to specific mixtures, we found no systematic evidence of mixture suppression or enhancement (as defined by neurons whose response profile would be fit with a V shape function). This discrepancy may be related to the possible inability of calcium imaging to resolve the brief suppressive patterns observed in the first 0.5 s following stimulus delivery and/or to our limited focus on superficial layers. A second, and more meaningful, difference from previous electrophysiological experiments in GC relates to the presence of a significant proportion of neurons that displayed categorical responses emerging in the delay period prior to the directional licking. Population decoding analysis highlighted differences in mixture coding between the sampling and delay periods. During the sampling period, classification performance increased linearly for progressively more distinct mixtures, with performance close to chance for the most similar mixture pairs (60/40 vs 40/60). On the contrary, the classification performance during the delay period was elevated for all pairs, including 60/40 vs 40/60. The confusion matrix for the decoding of all mixtures further confirmed these observations, showing virtually no mistakes for mixtures instructing different actions during the delay.

It is worth noting that the categorical responses described above could reflect the encoding of either the prevailing taste quality (sweet vs salty) or the predictive value of the mixture (lick left vs lick right). Analysis of error trials showed that the difference in responses to mixtures predicting different outcomes observed during the delay period was related to the upcoming action and not just to the prevailing taste quality.

Altogether, our data show a within-trial progression from linear, monotonic responses representing the concentration of individual components to the emergence of categorical responses encoding upcoming choice.

### Discrimination Learning and Taste Coding

Much of the existing work on discrimination learning has been performed in the visual, auditory, somatosensory, and olfactory systems^2,3,5-7,23,31^, and comparatively little is known about discrimination learning in taste. Studies on a variety of sensory systems have been instrumental in highlighting two general models accounting for the plasticity observed in discrimination learning^2-4,6,7,19^. According to one model discrimination learning is associated with the sharpening of sensory representations in sensory cortical areas^2,3,6,7^. A second model points at enhanced decision-related activity, with “higher-order” brain regions improving their ability to interpret sensory information to guide actions^8-11^. While these two perspectives are not necessarily incompatible, the focus on different brain regions (sensory vs high-order) has created a dichotomy. Recent studies in mice have challenged a hierarchical view of the brain and shown that decision-related responses necessary for making accurate choices can also emerge in primary sensory regions^17,23,31^. Using a taste discrimination task, our lab has recently shown the coexistence of sensory and decision-related signals in in GC^17^, opening a question as to what plasticity mechanism, if any, may be at play for GC in taste discrimination learning.

GC encodes sensory, affective and cognitive signals through time varying patterns of activity ^11,13,32-37^. Electrophysiological recordings in mice performing a 2-AC show a temporal separation between sensory processing and decision-related activity, with the first occurring mostly during sampling and the second manifesting itself as preparatory activity preceding decisions on directional licking^17^. This separation of signals, which was also observed in the present study with two photon calcium imaging, is advantageous for investigating plasticity associated with discrimination learning as it allows for the isolation of changes in sensory and decision-related activity.

Despite the advantages offered by GC as a model for studying plasticity^33,38,39^ and its well-demonstrated role in conditioned taste aversion^33,36^, familiarity learning^40,41^ and associative learning^32,42,43^, few studies have investigated GC’s involvement in taste discrimination learning. Work in rodents has shown that lesions of GC impact salt discrimination learning, confirming a role for this region in this process^44^. Here we developed a discrimination learning paradigm that is directly inspired by protocols used for other sensory modalities^18,19,23^ and that is amenable for imaging and electrophysiological studies in restraint mice. By testing mice with a battery of sucrose and NaCl mixtures and training them to discriminate progressively more similar pairs of stimuli, we demonstrated the effects of learning on taste processing. Imaging GC activity in the superficial layers revealed a series of changes associated with learning. The changes mostly occurred in the delay period, with an increase in the proportion of neurons selectively responding to stimuli predicting different licking directions and in the magnitude of selectivity. Similarly, we saw an increase in the neurons whose response profile was sigmoidal and an overall increase in categorical representation of choice selectivity. Altogether these results show that discrimination learning changes decision-related signals in GC, enhancing its ability to interpret gustatory information to drive decisions.

While we did not observe changes in taste processing, our experiments cannot rule out that spiking responses representing gustatory information in other layers of GC may have been enhanced. However, taking our results face value and assuming no plastic changes in sensory representations raises the question of how sensory responses can be transformed by GC circuitry to improve decision-making activity. A large body of evidence shows that GC activity is metastable, with ensembles going through transient patterns of coordinated spiking activity^35,45-50^. Each state can last up to a few seconds (∼100 ms to ∼3 s), before suddenly transitioning into another. Metastable states encoding for both sensory and decisional variables have been observed in GC of mice performing a 2-AC^49^. It is possible that taste learning may lead to plasticity of GC circuits such that there is improved ability to undergo network transitions from sensory to decision coding states. That is, while discrimination learning may not result in plasticity of single neuron sensory representation, it may affect network dynamics and create more reliable metastable sequences transitioning sensory signals into choices. Of course, the contribution of external sources in enhancing decision-related signals cannot and should not be ruled out. Future work using high density electrophysiological recordings to resolve metastable dynamics across various layers of GC in mice performing the taste mixture discrimination task may provide further insights on the neural mechanisms underlying taste discrimination learning.

In summary, this study used a novel taste mixture discrimination learning paradigm to i) investigate the coding of mixtures in GC and ii) demonstrate that decision-related responses are enhanced after learning and improved mixture discrimination performance.

## ACKNOWLEDGEMENTS

The authors would like to acknowledge Dr. Arianna Maffei, Dr. Memming Park, Ayesha Vermani, Dr. Joshua Plotkin, Dr. Anissa Abi-Dargham, Dr. Qiaojie Xiong, and past and present members of the Fontanini and Maffei laboratories for their feedback and insightful comments. This work has been supported by National Institutes of Health grants F30DC019523 to JFK, and UF1NS115779 and R01DC018227 to AF.

## AUTHOR CONTRIBUTIONS

Conceptualization, JFK and AF; Methodology, JFK and AF; Investigation, JFK; Formal Analysis, JFK; Writing – Original Draft, JFK and AF; Writing – Review and Editing, JFK and AF; Supervision, AF; Funding Acquisition, AF.

## DECLARATION OF INTERESTS

The authors declare no competing interests.

## METHODS

### Key Resources Table

| Reagent or Resource | Source | Identifier |
| --- | --- | --- |
| **Bacterial and Virus Strains** |  |  |
| AAV1-syn-jGCaMP7f-WPRE | Gift from Douglas Kim & GENIE Project | Addgene viral prep # 104488-AAV1, RRID:Addgene_104488 |
| **Experimental Models: Organisms/Strains** |  |  |
| Mouse; PV-cre | The Jackson Laboratory | JAX stock #017320, RRID:IMSR_JAX:017320 |
| Mouse; Ai14 | The Jackson Laboratory | JAX stock #007908, RRID:IMSR_JAX:007908 |
| **Software and Algorithms** |  |  |
| MATLAB | The MathWorks | https://www.mathworks.com/products/matlab.html, RRID:SCR_001622 |
| CalmAn | Github | https://github.com/flatironinstitute/CalmAn, RRID:SCR_021477 |
| GraphPad Prism 8 | GraphPad | https://www.graphpad.com/scientific-software/prism/, RRID:SCR_002798 |
| Fiji (ImageJ) | NIH | https://imagej.net/Fiji, RRID:SCR_002285 |
| LabView 2011 | National Instruments | https://www.ni.com/en-us/shop/labview.html, RRID:SCR_014325 |
| **Other** |  |  |
| Two photon microscope | Sutter Instruments | MOM, RRID:SCR_018860 |
| Microinjection syringe pump | World Precision Instruments | Cat #: UMP3T-1 |
| RHD2000 USB Interface Board | Intan Technologies | Cat #: C3100 |
| Micro-prism | Tower Optics | Cat #: MPCH-2.0 |
| NI-DAQ board | National Instruments | Cat #: USB-6001 |
| Motors | Zaber Technologies | Cat #: X-LSM025B |
| Solenoid valves | The Lee Company | Cat #: LHDA123115H |

### Resource Availability

#### Lead Contact

Further information and requests for resources and reagents should be directed to and will be fulfilled by the Lead Contact, Alfredo Fontanini.

### Materials Availability

This study did not generate new unique reagents.

### Data and Code Availability

Datasets and code supporting the current study are available upon request from Lead Contact.

### Experimental Model and Subject Details

Adult male and female mice were used (8 to 12 weeks at time of surgery). The mice used were transgenic PV-cre;Ai14 mice heterozygous for both Cre and Lox-Stop-Lox-tdTomato alleles and were bred in house by crossing female homozygous PV-cre^51^ (B6.129P2-*Pvalb^tm1(cre)Arbr^*/J, JAX stock #017320) mice with males homozygous for the Ai14 reporter gene^52^ (B6;129S6-*Gt(ROSA)26Sor^tm14(CAG-tdTomato)Hze^*/J, JAX stock #007908). These animals were initially chosen to isolate signal from PV+ inhibitory neurons, however due to the relatively low yield of PV+ cells in GC, all recorded neurons were pooled for analysis. Mice were maintained on a 12 hour light/dark cycle with *ad libitum* access to food and water unless otherwise specified. Mice were group housed until prism implant surgery and then single housed. All experimental protocols were approved by the Institutional Animal Care and Use Committee at Stony Brook University, and complied with university, state, and federal regulations on the care and use of laboratory animals.

### Surgical Procedures for Viral Injection and Prism Implantation

Surgeries were performed as previously described^13,38,42^. Mice were anesthetized with a cocktail of ketamine (70 mg/kg) and dexmedetomidine (1 mg/kg) administered via intraperitoneal injection. Bupivacaine (2 mg/kg) was administered under the scalp for local anesthesia, and carprofen (5 mg/kg) and penicillin (150000 UI/kg) were administered via subcutaneous injection for analgesia and infection prevention respectively. Dexamethasone was also injected subcutaneously to reduce brain swelling (0.2 mg/kg) and ophthalmic ointment was applied to the eyes to prevent drying. Depth of anesthesia was assessed by checking toe pinch reflex and body temperature was maintained at 37°C using a heating pad. Mice were placed in a stereotaxic apparatus and a midline incision made in the scalp. The skull was leveled, and a craniotomy performed above GC. Virus expressing gCaMP7f (AAV1-syn-jGCaMP7f-WPRE, stock concentration 3 x 10^13^ vg/mL, Addgene, 104488-AAV1)^53^, was loaded into a Hamilton syringe and injected at using a microinjection syringe pump (World Precision Instruments, UMP3T-1) at two AP locations (+1.1 mm and +0.8 mm anterior, ∼3.7 mm left from bregma, close to the lateral suture). At each location, injections were performed at two DV sites (1.6 mm and 1.9 mm from pial surface). Virus was infused at 1 nl/s with a volume of 200 nl/site (total volume 800 nl across four sites). The pipette was lowered to the first DV site left in place for 5 minutes after the injection was finished, and then lowered to the second site and left in place for 15 minutes before being slowly retracted. After injections were complete, the scalp was closed using surgical sutures and cyanoacrylate glue (3M, Vetbond). After surgery was complete, Antisedan (atipamezole hydrochloride, 1 mg/kg) and lactated ringers solution were administered subcutaneously to reverse anesthesia and for hydration. Mice were allowed to recover for 1 - 2 weeks before the prism implantation and headpost surgery.

Microprisms were fabricated as previously described^13,38,42^. Mice were anesthetized and placed in the stereotaxic apparatus, the scalp and skin overlying the left temporalis muscle was removed. The left temporalis muscle was then dissected to expose the skull overlying GC, and a ∼2.2 x 2.2 mm craniotomy was performed on the lateral surface of the skull overlying GC using a dental drill. The craniotomy was cleaned, and a durotomy was performed using fine forceps. The microprism was carefully lowered into the craniotomy with the ventral edge inserted below the zygomatic bone and sealed in place with cyanoacrylate. The skull was covered with a layer of UV curable dental adhesive (Henry Schein, iBond self-etch), and the prism and a custom headpost were secured with dental cement. Carprofen was administered postoperatively for five days for analgesia and to reduce inflammation.

### Taste Mixture Discrimination Paradigm

At least one week after prism surgery, mice were placed on water restriction (target 85% body weight, ∼1.5 ml per day) for one week before starting training. The training was performed in several phases to habituate the mice to the restraint apparatus and measure neural activity and performance across learning. First, mice were habituated to head fixation and trained to lick a drop of water from the motorized central spout (1 - 2 days). Next, they were habituated to the trial structure. Mice had to learn to lick a drop of water from the central spout. After they licked at least once, the spout retracted, and the lateral spouts advanced following a ∼3.5 s delay. A drop of water (5 µl) was released after one lick to whichever side was licked first. Once they were consistently licking the central and lateral spouts (usually 1 - 3 days), the mice began pre-training on the mixture discrimination. A preformed drop (4 µl) of either sucrose (100 mM) or NaCl (100 mM) was presented at the central spout, and mice had to lick once on the correct side to receive a drop of water (5 µl). Correct sides for sucrose and NaCl choices were counterbalanced across animals, and there was no difference in the latency to left versus right lateral licks (paired t test, p = 0.66). Errors were punished with no water and a five second timeout. After reaching criteria performance in the pre-learning phase (85% performance for three consecutive days, see data analysis section for calculations of performance), mice were given a pre-learning psychometric test over two consecutive sessions. They were given six different stimuli presented in a pseudorandom order, and mice were rewarded for a correct choice if they licked towards the direction indicating the predominant taste in the mixture. The stimuli were (%sucrose/%NaCl): Day 1: 100/0, 85/15, 65/35, 35/65, 15/85, 0/100; Day 2: 100/0, 75/25, 60/40, 40/60, 25/75, 0/100. Next, the mice entered the discrimination learning phase, where they were trained on increasingly similar mixture pairs. First on 75/25 vs 25/75, then 65/35 vs 35/65, and finally 60/40 vs 40/60. For the first two pairs, criteria performance was 85% correct for three consecutive days up to a maximum of seven total sessions and for the 60/40 vs 40/60 sessions, the criteria performance was 80% correct for three consecutive days. Finally, the animals were tested on a post-learning psychometric test, with the same stimuli as the pre-learning test.

### Taste and Water Delivery Apparatus

The central and lateral spouts, as well as a vacuum line were mounted on three separate motors (Zaber, X-LSM025B). After each trial, a small drop of water was dispensed on the central spout and cleared by vacuum aspiration to ensure no residual taste would be present for the following trial. Taste solutions were made fresh each day and were dispensed from gravity fed solenoid valves (The Lee Company, LHDA123115H). Valves were commanded by valve controllers (Automate Scientific, ValveLink 8.2) driven through NI-DAQ boards (National Instruments, USB-6001) by custom LabVIEW scripts. All events and imaging frames were synchronized and recorded on an Intan RHD2000 device.

### Two Photon Imaging

Imaging was performed on a commercial two photon microscope (Sutter MOM) using a resonant scanning module controlled by MScan software (Sutter) mounted with a 10x super apochromatic objective (Thorlabs, 0.5 NA, 7.77 mm WD). Two photon excitation of gCaMP7f was accomplished using a Ti:Sapphire femtosecond laser (Coherent) tuned to 980 nm with a laser power of 100-250 mW at the front of the objective and emission was collected using photomultiplier tubes (Hamamatsu). Images were acquired at 31 Hz and triggered two seconds before the start of each trial (trial start was defined as when the central spout begins to advance).

### Histological Staining

Mice were deeply anesthetized and transcardially perfused with phosphate-buffered saline (PBS), followed by perfusion with 4% paraformaldehyde (PFA) in PBS. Brains were dissected out and postfixed in 4% PFA at 4°C for 24 hours. Thin (50 µm) coronal brain sections containing the GC were cut on a vibratome (Leica, VT1000). Brain sections were first washed in PBS (3 × 5 minutes) at room temperature (RT), then incubated with the nuclear stain, Hoechst 33342 (1:5000, Invitrogen, H3570) in PBS for 30 minutes at RT. Sections were then washed with PBS (3 x 10 minutes) and mounted onto glass slides with Fluoromount-G. All sections containing gCaMP7f signal were mounted and imaged on a fluorescence microscope (Olympus BX51WI) at 4x magnification.

### Data Analysis and Statistics

Data analysis was performed using custom scripts written in MATLAB 2019a (MathWorks), ImageJ (NIH), and Prism 9 (GraphPad).

### Behavioral Analysis

Behavioral performance was calculated by dividing the total number of correct trials by the total number of trials performed. Trials where mice did not lick to the central or lateral spout were excluded, typically these only occurred at the end of a behavioral session. Psychometric curves were calculated by fitting a sigmoid function to the mean performance for each of the tastants. Psychometric curves were compared between the pre- and post-learning groups using an extra-sum-of-squares F test to test if one or two curves were a better fit to the dataset. Mixture performance curves were calculated using the mean performance for each mixture pair, and differences in mixture performance were compared within the pre-learning group by one way ANOVA. Mixture pair performance was compared by fitting the data with a logistic function and using an extra sum-of-squares F test to test if one or two curves were a better fit to the dataset.

### Two Photon Image Processing

Calcium activity from each trial were saved as individual multiframe TIFF files, which were down sampled from 31 Hz to 6 Hz using ImageJ (grouped Z project) and concatenated for each session. Rigid motion correction was applied using the NormCorre algorithm^54^ and regions of interest (ROIs) corresponding to individual neurons were extracted using the CNMF algorithm^55^. ROIs and ΔF/f traces were manually inspected and curated using a custom GUI built in MATLAB (MathWorks) and ΔF/f values were used for all downstream data analysis.

### Neural Data Analysis

All analysis of neural data was performed on the second psychometric testing session when the following stimuli were presented (%sucrose/%NaCl): 100/0, 75/25, 60/40, 40/60, 25/75, 0/100. We did not attempt to track neurons across sessions, thus could not compare responses from the same neurons across multiple psychometric testing sessions.

Responsive neurons were identified using only correct trials, grouped into sucrose and NaCl predominant trial types. Sampling responsive neurons were identified by aligning data to the first lick to the central spout (onset of sampling) and comparing the mean activity between the 1.5 s window after sampling onset to a pre-stimulus baseline window (-4 s to -2.5 s before sampling onset, rank sum test, p < 0.05). Delay responsive neurons were similarly identified by aligning the neural activity to the first lick to the lateral spout (choice onset) and comparing a 1.5 s window before the choice to a pre-stimulus baseline window (-6.5 s to -5 s before choice, rank sum test, p < 0.05). Selective neurons were identified using this subset of sampling and delay responsive neurons by comparing activity between the two trial types in the sampling or delay periods (rank sum test, p < 0.05). Single neuron average responses were calculated by grouping correct trials by predominant taste in the mixture to obtain two trial types (sucrose and NaCl predominant). Neural activity was aligned to either the onset of sampling or choice, averaged across all trials and plotted as mean ± SEM. Population average responses were plotted by averaging correct trials across all trial types for each of the sampling or delay responsive neurons, and then finding the mean ± SEM across all neurons.

Curve fitting analysis was performed as previously described^21^. Neural activity was aligned to sampling and choice onset and binned in consecutive 0.5 s bins. Mean activity for all responsive neurons in each of the six trial types was calculated for each bin and fit with both a linear and sigmoid function. Neurons were assigned as having a best fit to either the linear, sigmoid or neither function based on a neuron having a significant fit (p < 0.05) and the largest F value. This approach accounts for the larger number of parameters required to fit the sigmoid function. Proportions of linear and sigmoid best fits were compared using a χ^2^ test (p < 0.05).

Population decoding was performed as previously described using the maximum correlation decoder^17,25^. Data from all neurons and all taste mixtures were combined into a pseudopopulation with six class labels corresponding to the six taste stimuli. Only data from correct trials were included in the analysis. The mean responses in the sampling (0 s to 1.5 s after sampling onset) and delay (-1.5 s to 0 s before choice) windows were calculated for each trial. Data were randomly divided into ten splits, nine were used for training and the remaining used for testing the decoder. This process was repeated ten times with different testing/training splits, and decoding accuracy across all testing splits was used to calculate overall decoding performance (tenfold cross validation). For decoding of all six mixture stimuli, confusion matrices were also generated to assess errors in the decoder. For calculation of decoding performance in mixture pairs, separate decoders were trained for each of the three mixture pairs (100 vs 0, 75 vs 25 and 60 vs 40). Overall decoding performance was calculated as the percentage of trials where the correct taste mixture was classified using tenfold cross validation.

To calculate stimulus and choice selectivity, trials were again grouped into two trial types based on the predominant taste in the mixture, and neural activity was separated into sampling and delay periods. Stimulus selectivity was calculated by finding the difference in mean activity between trial types on correct trials only. This value will be positive if a neuron is more responsive to sucrose and negative if it is more responsive to NaCl. Choice selectivity was calculated by finding the difference in mean activity between trials with the same stimulus but a different choice (correct vs error trials). This value was calculated separately for sucrose and NaCl trials and then averaged for each neuron. A neuron with a high choice selectivity will differentiate between left and right choice, even when the taste delivered is the same. This value will be positive if a neuron responds to the direction associated with sucrose and negative if it responds to the direction associated with NaCl. The actual direction (left or right) can vary because the port associated with each stimulus was counterbalanced across animals. Stimulus vs choice selectivity was plotted and r^2^ values and line slopes were calculated (significant if p < 0.05).

Comparisons between pre- and post-learning were done using a χ^2^ test (p < 0.05). Mean stimulus selectivity (correct trials only) was compared between pre- and post-learning using an unpaired t test (p < 0.05). Proportions of neurons with a significant linear or sigmoid fit were compared using a χ^2^ test (p < 0.05). Category tuning index was calculated as described previously^9,10^, and compared using an unpaired t test (p < 0.05).

### Equations

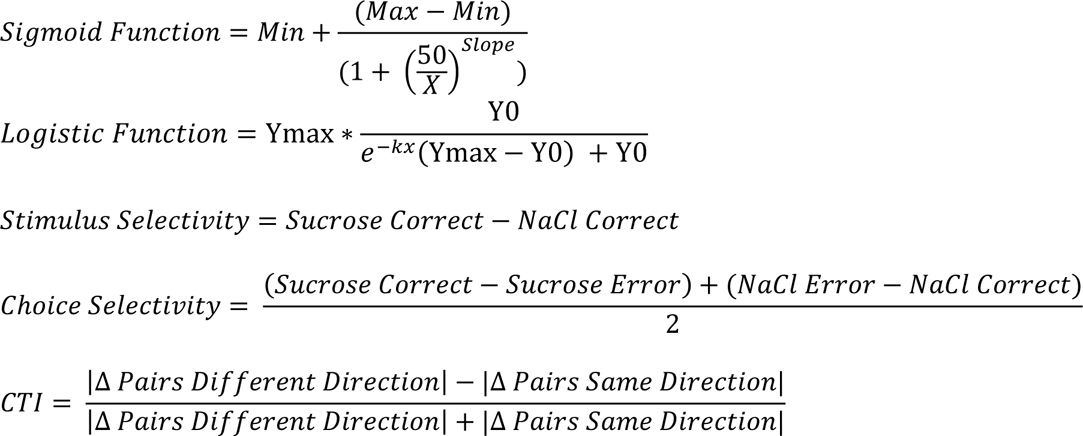

## FIGURES

**Supplemental Figure 1.**
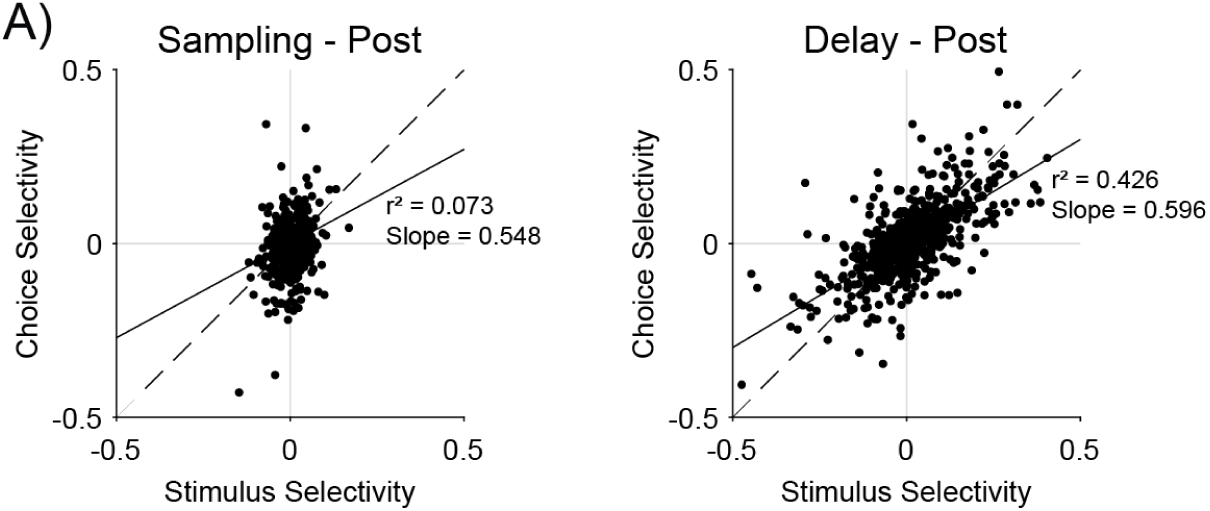
Post-Learning Delay Period Activity and Coding of Upcoming Choice. **A)** Scatter plots showing stimulus versus choice selectivity for sampling (left) and delay (right) responsive neurons in the post-learning condition. The solid line represents a linear fit to the data (sampling: r^2^ = 0.073, slope = 0.548, p < 0.001; delay: r^2^ = 0.426, slope = 0.596, p < 0.001) and the dashed line represents the unity line.

## Notes

### Competing Interest Statement

The authors have declared no competing interest.

